# Cryo-EM structure of the Type IV pilus extension ATPase from enteropathogenic *Escherichia coli*

**DOI:** 10.1101/2022.08.10.503557

**Authors:** Ashok R. Nayak, Pradip K. Singh, Jinlei Zhao, Montserrat Samsó, Michael S. Donnenberg

**Author notes:** Corresponding authors*. Email addresses.

## Abstract

Type 4 pili (T4P) are retractable surface appendages found on numerous bacteria and archaea that play essential roles in various microbial functions, including host colonization by pathogens. An ATPase is required for T4P extension, but the mechanism by which chemical energy is transduced to mechanical energy for pilus extension has not been elucidated. Here we report the cryo-electron microscopy (cryo-EM) structure of the BfpD ATPase from enteropathogenic *Escherichia coli* (EPEC) in the presence of either ADP or a mixture of ADP and AMP-PNP. Both structures, solved at 3 Å resolution, reveal the typical toroid shape of AAA+ ATPases and unambiguous six-fold symmetry. This six-fold symmetry contrasts with the two-fold symmetry previously reported for other T4P extension ATPase structures, all of which were from thermophiles and solved by crystallography. In the presence of the nucleotide mixture, BfpD bound exclusively AMP-PNP and this binding resulted in a modest outward expansion in comparison to the structure in the presence of ADP, suggesting a concerted model for hydrolysis. *De novo* molecular models reveal a partially open configuration of all subunits where the nucleotide binding site may not be optimally positioned for catalysis. ATPase functional studies reveal modest activity similar to that of other extension ATPases, while calculations indicate that this activity is insufficient to power pilus extension. Our results reveal that, despite similarities in primary sequence and tertiary structure, T4P extension ATPases exhibit divergent quaternary configurations. Our data raise new possibilities regarding the mechanism by which T4P extension ATPases power pilus formation.

## INTRODUCTION

Type 4 pili (T4P) are the most ancient and widespread class of pili, produced by Gram-positive and Gram-negative bacteria and by archaea (1). T4P are composed of thousands of copies of a major pilin protein arranged in a helical array (2). They are assembled and retracted by a complex machine composed of several essential proteins, including a dedicated extension ATPase most often called PilB that is essential for pilus biogenesis. Prior to incorporation into the pilus, pilin is an integral transmembrane protein (3). Thus, PilB is thought to provide mechanical energy required to extricate the pilin from the cytoplasmic membrane to assemble the pilus. The enzyme activities of several purified T4P extension ATPases have been measured *in vitro*, with reported rates varying from 0.7 to 700 nanomoles of ATP hydrolyzed per minute per mg protein (4-9). How this *in vitro* activity relates to pilus extension, which has been measured in microns per second (10-12), remains unclear. Many bacteria also have one or more dedicated retraction ATPases (4, 10, 12, 13).

Precisely how conformational changes induced by ATP hydrolysis are transduced by PilB to lift and extricate pilin from the membrane so that it can be added to the base of the growing pilus remains an unanswered question (14). High-resolution structures of catalytic domains of three closely related PilB-family ATPases solved by X-ray crystallography revealed a two-fold symmetry of the hexamers (14-17), whereas other members of the AAA+ family of ATPases display six-fold symmetry (18-20). The PilB structures, all from thermophilic Gram-negative bacteria, have several other features in common: (1) all are hexamers; (2) the first N-terminal domain (N1D) is not visualized; (3) a flexible linker separates the second N-terminal domain (N2D) from the C-terminal domain (CTD), which contains the catalytic site; (4), a density is consistent with a Mg^2+^ ion and (5) remote from the active site, is a zinc-binding motif. Within each structure are three pairs of subunits on opposite sides of the hexamer. In one pair, relative to the other two, the N2D is rotated towards the center of the toroid (NTD-in), while in the other two pairs the N2D is rotated away from the center (NTD-out). A symmetric, rotatory mechanism of hydrolysis has been proposed, which results in a “scooping” motion in which the CTD is displaced upward and toward the center of the ring, where it could translate this motion to membrane-bound pilin (14, 15).

PilB from *Thermus thermophilus* (TtPilB) was also examined by cryo-EM, both bound to the non-hydrolysable ATP analogue adenylyl-imidodiphosphate (AMP-PNP) and without exogenous nucleotide. Although resolution of only ∼8 Å was achieved, this structure showed for the first time the second and third of three predicted N1Ds (16). This more complete structure showed two hexamers joined by a constriction. One of the hexamers, into which the N2D-CTD crystal structure was docked, showed clear two-fold symmetry. The other hexamer, presumably representing the second and third N1Ds, appeared to display six-fold rather than two-fold symmetry. Comparison of the hexamers in the presence of AMP-PNP and without added nucleotide showed little evidence for the symmetric rotary model, nor for translation of movement through the center of the multimer. Instead, cryo-EM shows an outward shift in the center of mass of the AMP-PNP structure relative to the structure solved without addition of nucleotide, rather than the change in the orientation of the N2D-CTD protomers seen by crystallography (14, 15). Evidence of displacement of the N1D hexamer by 10-13 Å was also reported. The authors suggested an alternative model linking the N1D displacement to pilin extrication.

Overall, while available PilB structures have provided valuable information, there is no agreement yet on the significance of the two-fold symmetry, whether it is critical to explain the mechanism by which chemical energy is converted to mechanical energy, and which structural changes are caused by ATP hydrolysis.

Here, we focused on the extension ATPase from a Gram-negative human pathogen, enteropathogenic *Escherichia coli* (EPEC), which expresses a bundle-forming pilus (BFP) distantly related to T4P of thermophiles (21). We purified the full-length EPEC PilB homologue, BfpD, determined its structure by cryo-EM both in the presence of ADP alone and in the presence of a mixture of ADP, ATP, and AMP-PNP, achieving unprecedented resolutions of 3.0 and 3.1 Å, respectively, and measured its enzyme activity. The six-fold symmetry that we observed suggests a concerted, rather than a symmetric rotary mechanism of energy coupling that may have implications relevant to all PilB family members.

## RESULTS

### Negative staining and cryo-EM of BfpD reveal a six-fold symmetry

Purified BfpD was prepared under two nucleotide conditions: either in the presence of ADP (BfpD-ADP dataset), or in the presence of ADP, ATP and the non-hydrolysable ATP analog, AMP-PNP (termed BfpD-ANP dataset) at a ratio of 2:4:5. We reasoned that the presence of ATP would allow completion of catalysis culminating in occupancy by the nucleotide preferred by each subunit, depending on its position in the catalytic cycle. Examination of BfpD-ANP by negative staining in the presence of a reducing agent showed the expected toroid structure, and further reference-free alignment showed 2D averages with six-fold symmetry with protruding edges (Fig. S1).

After confirming structural integrity of the sample, cryo-EM and high throughput data collection yielded 7,207 good movies and 313,223 good particles for BfpD-ADP. For BfpD-ANP, a dataset of 9,853 good movies and 424,708 good particles with well-distributed orientation yielded two classes, with 214,673 and 58,944 particles, respectively. Reference-free 2D averaging demonstrates unambiguous six-fold symmetry in both nucleotide states (Fig. S2-S3). Subsequent 3D reconstruction with C6 symmetry yielded resolutions of 3.1 Å, 3.0 Å and 3.7 Å for BfpD-ADP, BfpD-ANP class-1 and BfpD-ANP class-2, respectively (Figs.1, S2-S3). BfpD-ANP class-1 has an outward N2D compared to BfpD-ANP class-2. The dimensions of BfpD are 130 Å maximum width, 68 Å height, with a central pore measuring 30 Å in diameter on one side and twice this diameter on the other.

### Overall BfpD structure, active site, and hexameric association

*De novo* models were built for residues 107-219, which represents the N2D and 232-534, which represents the CTD (Fig. 2A, B). N1D and the eight residues intervening the two domains were not visualized, probably owing to intrinsic flexibility, as has been reported for other T4P extension ATPases (14-17). The catalytic site is formed by the interface between the Arginine finger (Arg 217) of N2D, the Walker A motif (260-267), and the catalytic loop of the CTD (294-310) of each subunit (Figs 2-3). For the BfpD-ANP dataset, despite the addition of ADP, ATP, and AMP-PNP, the active site of all monomers was fully occupied by electron density consistent with AMP-PNP, suggesting that ultimately ANP occupied the active sites and remained there. The presence of ANP in the catalytic site results in an outward N2D movement, stabilized by interaction between Arg 217 and the γ-phosphate. Both structures show catalytic glutamate residue Glu295 and Walker B motif Glu338, 5.3-11.6 Å and 6 Å, respectively, from the phosphorus atom in the terminal nucleotide phosphate (Figs. 2, 3). The loop (294-310) carrying the catalytic Glu295 has a well-defined cryo-EM density and is positioned closer to the nucleotide in BfpD-ADP compared to BfpD-ANP, assisted by a hydrogen bond between ADP and Tyr296 (Figs. 2, 3). Additionally, in the BfpD-ADP map, Mg^2+^ is seen between Ser267 (from the Walker A motif) and the beta-phosphate of ADP. As in other PilB-family members, Zn^2+^ is observed far from the catalytic site and from the subunit interface (Fig. 2B), coordinated by the zinc-finger-like tetracysteine motif (Cys403, Cys406, Cys445, Cys446).

The electrostatic map shows the toroid surface lined by the N2D domains as the side that likely faces the membrane, because, in comparison to the CTD side, its positive charge density is significantly higher (Fig. 4 A, B). Each subunit forms a chevron-like structure pointing to the right (+ subunit) when seen sideways with the N2D on top (Figs. 1, 4C). At the inter-subunit interface, the outer corner of the chevron, formed by the CTD, points towards the inner corner of the neighboring (+) subunit, interacting by the N2D/CTD and CTD/CTD+ interfaces, with ∼1400 Å^2^ buried surface area. The ATP binding site is not involved in the inter-subunit interaction. Several residues face each other within 3.8 Å, suggesting potential salt bridges or hydrogen bonds between adjacent subunits. Starting from the N’ terminus of the subunit in the left, the pairs of residues likely forming hydrogen bonds are R328-Q203, N286-Y213, N286-K129, and those likely forming salt bridges are D330-R140, D332-R140, K383-D376, D386-R475 (Fig. 4C), where the second residue is from the (+) subunit.

**Figure 1.**
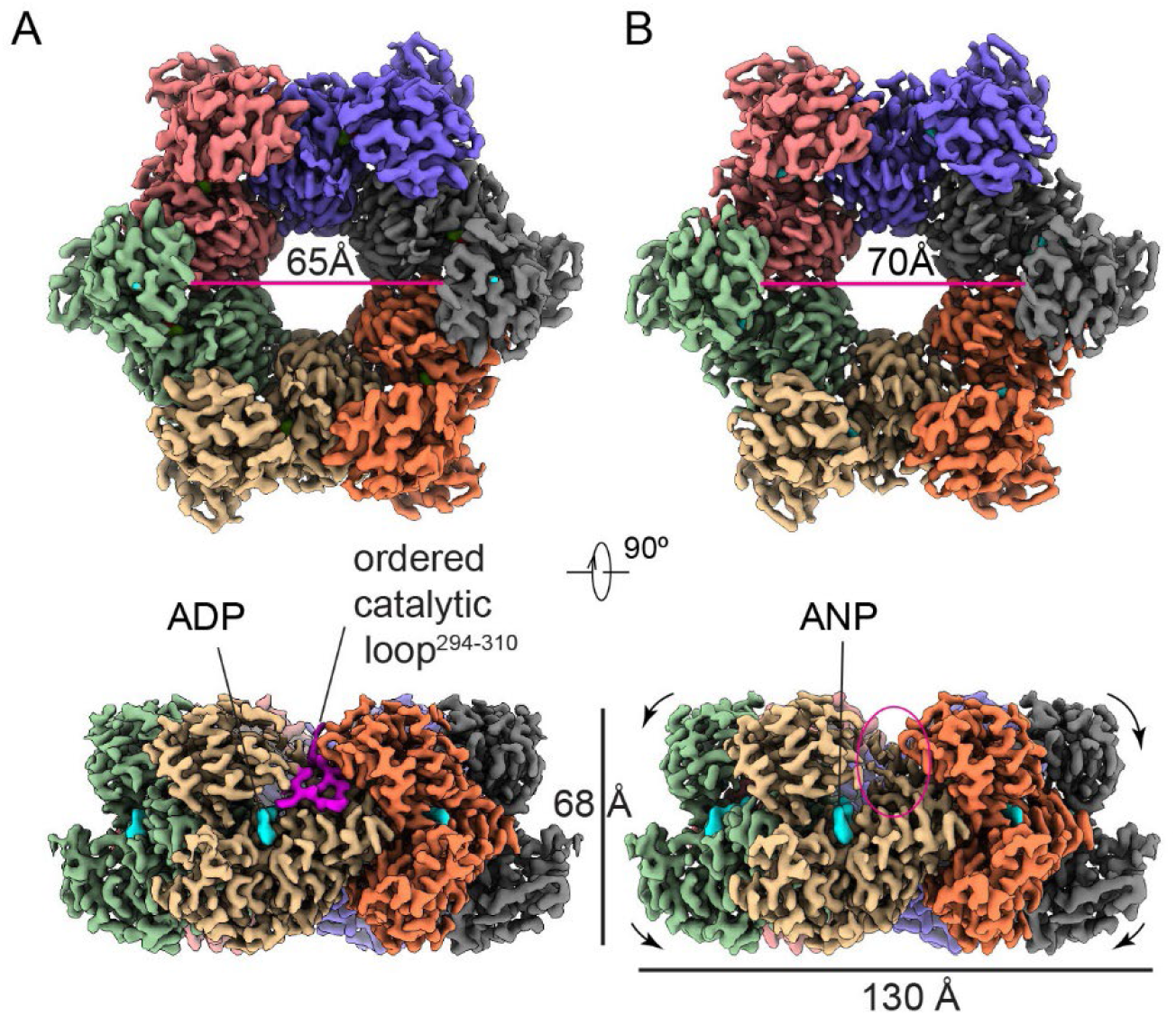
Cryo-EM maps of six-fold symmetric BfpD and its nucleotide binding pocket. 3D reconstruction of BfpD in (A) ADP and (B) a mixture of ADP, ATP, and ANP resolved at 3.0 Å and 3.1 Å resolution shown in two orthogonal orientations. Top row, view N2D facing the viewer. Bottom row, side view with N2D at the top. The diameter of BfpD toroid center expands by ∼5 Å (arrows) in the presence of ANP compared to the diphosphate. Each subunit is colored differently, and the nucleotide is represented in cyan. The EM density for the loop harboring catalytic Glu295 (magenta) is well defined in the ADP structure, but not in the ANP structure (red oval).

**Figure 2.**
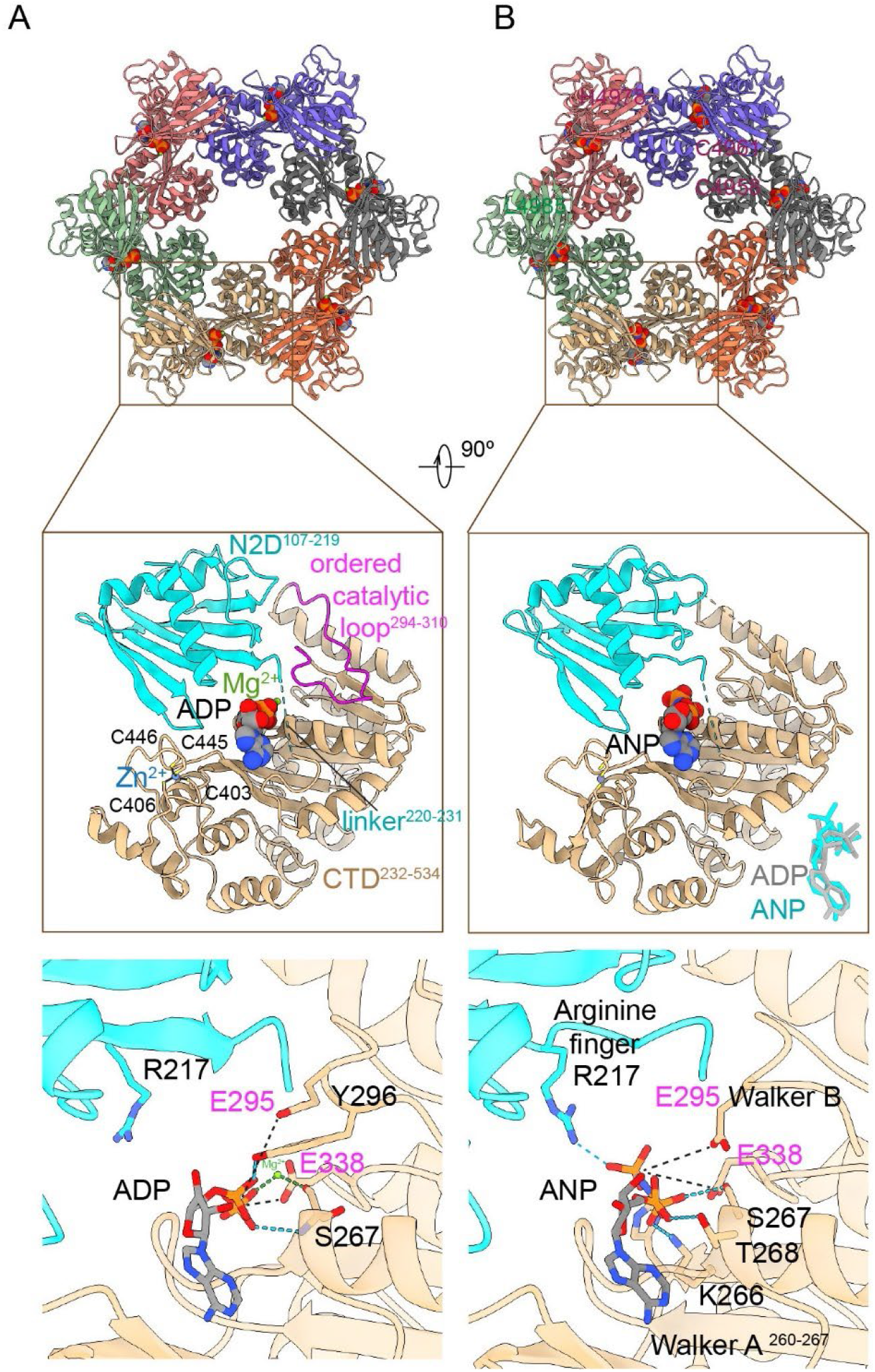
*De novo* models of BfpD-ADP (A) and BfpD-ANP (B). In the upper panel, each monomer is represented in a different color. The middle panel shows the domain organization of one subunit with the nucleotide, Mg^2+^, ordered catalytic loop and tetracysteine motif shown in BfpD-ADP. Key amino acids at the N2D-CTD domain interface responsible for nucleotide binding (black) and hydrolysis (magenta) are shown in the lower panel. The catalytic Glu295 and Walker B Glu338 are positioned at 5.3-11.6 and ∼6 Å in ADP and ANP models respectively. In the BfpD-ANP model, an interaction between the γ-phosphate and Arg217 repositions the loop carrying the catalytic Glu295.

**Figure 3.**
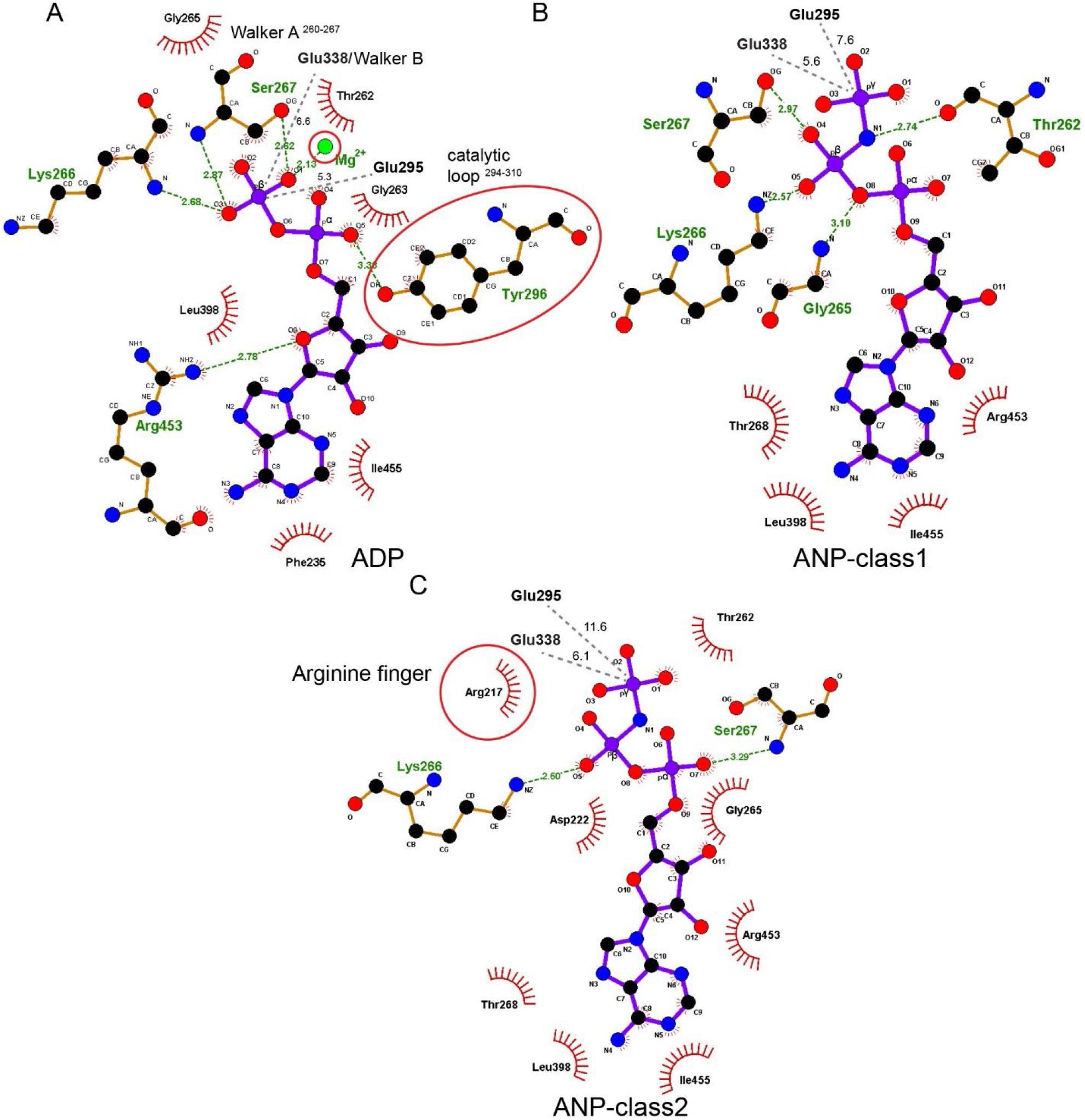
BfpD active site reorganization upon ATP hydrolysis. 2D representation of the BfpD catalytic site in (A) BfpD-ADP, (B) BfpD-ANP-class1, and (C) BfpD-ANP-class2 structures. The detail shows changes in the interaction network associated with the conformational change that occurs upon transition from dinucleotide to trinucleotide bound. Arg217 repositions to stabilize and place the γ-phosphate of ANP at a farther distance from Glu295. Hydrogen bonds, in Å, are shown with green dashed lines, non-bonded interactions are shown by red arcs. Tyr296 and Arg217 are highlighted to emphasize their important role in the indicated conformations.

**Figure 4.**
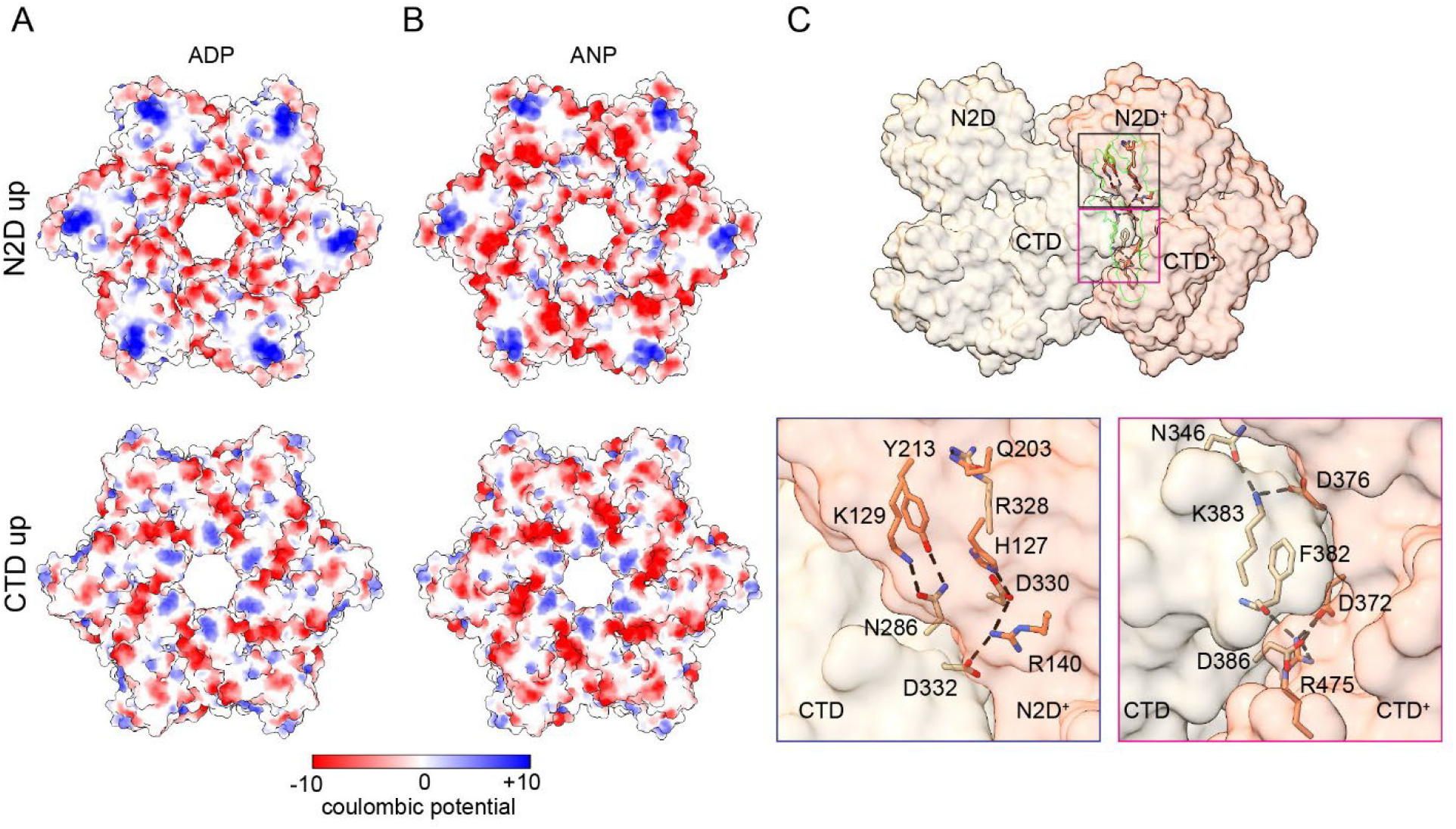
Charge distribution and interacting surfaces of BfpD. Surface electrostatics representation for BfpD-ADP (A) and BfpD-ANP (B). Top, the N2D facing surface of BfpD that has clusters of positively charged residues (in blue), suggesting this side most likely faces the plasma membrane. Bottom, CTD facing surface showing overall negatively charged residues (in red). (C) Top, side view of two neighboring BfpD subunits with the interacting space highlighted (green). Bottom, the CTD of the left subunit (tan) interacts with the N2D+ and CTD+ of the subunit (salmon) on the right. Salt bridges and hydrogen bonds <3.7 Å are shown.

### Presence of trinucleotide induces an expansion in the AAA+ ATPase while preserving the six-fold symmetry

Comparison of the BfpD-ADP and BfpD-ANP structures reveals a subtle outward shift in position of the N2D with respect to the CTD such that the mobile innermost N2D loops delineate a circle of 65 Å in diameter (distance between diagonally located Ala172 residues) in the presence of ADP, and 70-73 Å diameter in the presence of ANP class 1 and 2, respectively (Fig. 1). This change is accompanied by a slight change in angle of 1.1° for the N2D and 1.6° for the CTD between each monomer and the vertical axis for the BfpD-ANP structure compared to the BfpD-ADP structure (Movie S1), which is consistent with the outward shift in the center of mass described in the much lower resolution TtPilB cryo-EM structure (16). The conservation of six-fold symmetry in both BfpD structures suggests that the subunits may work in a concerted manner to translate chemical to mechanical energy.

### The N2D-CTD domains of BfpD have an intermediate rotation state

We compared BfpD with the four other T4P extension ATPases homologues, each of which has two-fold symmetry, to try to understand the mechanistic basis of force generation.

The TtPilB-ATPγS crystal structure (PDB ID:5IT5) is an elongated hexamer with two-fold symmetry (15). In TtPilB, all six subunits are bound to ATPγS. However, one pair of opposing subunits in the hexameric ring is in the N2D-in, or “closed” conformation, and the other four subunits are in the N2D-out “open” conformation. The four open N2D-out subunits are superimposable, as are the two closed N2D-in subunits (Fig. 5A). With the C-terminal domains superimposed, we found that the open subunits have a 57° outward rotation of their N2D domain, *i*.*e*., away from the center of the toroid (N2D-out), when compared to the closed subunits (Fig. 5A). Comparing BfpD-ANP to TtPilB-ATP indicated an intermediate conformation: the BfpD N2D is rotated by ∼42° from the TtPilB closed conformation and 18° from the TtPilB open conformation (Fig. 5B). Similar results were obtained for BfpD-ADP, as the rotation of N2D between the two nucleotide states was ∼1°.

**Figure 5.**
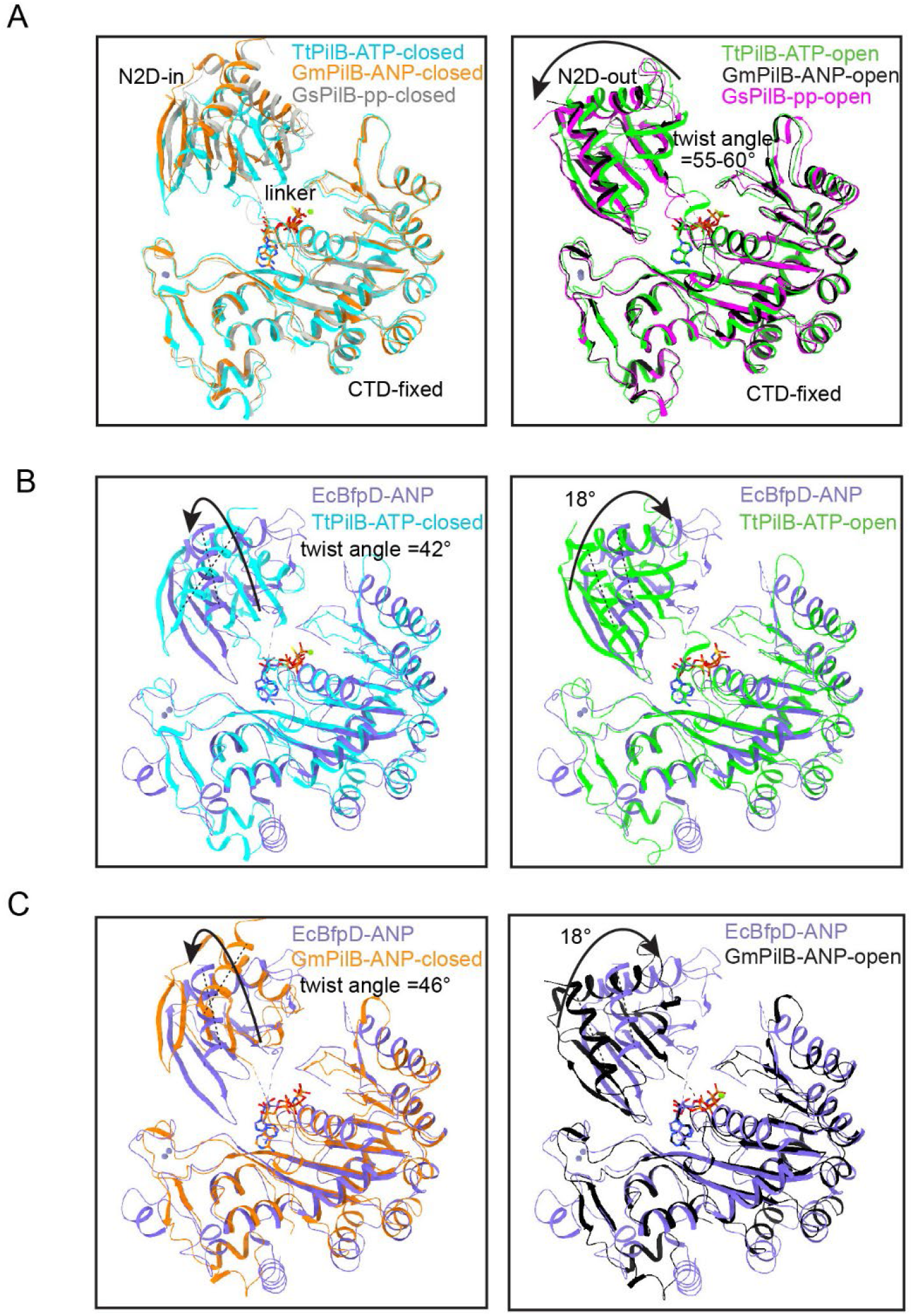
The N-terminal domain of BfpD has a distinct twist and rotation axis. (A) Two distinct conformations of pilus extension ATPase subunits were found in two-fold symmetric TtPilB, GmPilB, and GsPilB bound to ATP, AMP-PNP (ANP), and phosphate (pp) respectively. In A, the closed, left, and open, right, subunits are overlaid, keeping the CTDs fixed. The arrow shows rotation in N2D from closed to open conformation in each structure. In (B) and (C), the N2D domain of enteropathogenic *E. coli* BfpD is compared with the TtPilB-ATP and GmPilB-ANP structures, respectively. The arrow in the left indicates transition from closed TtPilB-ATP or GmPilB-ANP to BfpD-ANP. The arrow in the right indicates transition from TtPilB-ATP-open to BfpD-ANP. For the two-fold structures, the comparisons between BfpD and “open” are shown for the more extreme open (N2D-out) conformation; see Fig. S4.

The *Geobacter metallireducens* PilB (GmPilB)-ANP and GmPilB-ADP crystal structures are also two-fold symmetric hexamers, wherein four subunits are bound to ANP or ADP, respectively (14). The other two subunits in GmPilB-ANP and GmPilB-ADP structures bind to ADP and are empty, respectively. Two fully closed (N2D-in), two fully open (N2D-out), and two open intermediate subunits were found in GmPilB-ANP (PDB ID: 5TSH). Within each of the three pairs, the two subunits are superimposable (Fig. S4). The fully open pair of subunits are rotated ∼60° in their N2D from the closed subunits. However, the N2Ds of the intermediate-open subunits are rotated by 11° inward from the fully open subunits or rotated by 49° from the closed pair of subunits. BfpD-ANP showed ∼46° rotation in its N2D domain from the GmPilB-ANP closed subunits. Consistent with the three distinct kinds of subunits, the nucleotide-binding pockets of GmPilB at the N2D-CTD domain interfaces have different occupancy by ATP and thus varied affinity to the nucleotide. The N2D-in subunits have 100% occupancy of ANP and are proposed to be the active ATP hydrolysis interface. The catalytic Glu395 at this interface is located closest (at ∼6 Å) from the phosphorous atom of the ANP terminal phosphate. The other two pairs of subunits with ANP and ADP form an open and closed ATP binding pockets with N2D twists and 0° and 49° respectively and partial occupancy of 58-60% ANP and 44-54 % ADP, respectively. Larger γ-P distances of terminal phosphates, ∼10 Å at the ADP and 7-8 Å at the ANP interface, are suggestive of ATP binding and ADP/ATP exchange sites.

Furthermore, the *G. sulfurreducens* PilB (GsPilB) apo structure (PDB ID: 5ZFR) was also found to possess three distinct N2D conformations (17). Two pairs of the open N2D subunits have 66° and 59° rotations from the closed (N2D-in) pair of subunits (Fig. 5A). BfpD-ANP showed a ∼42° rotation in its N2D domain from the GsPilB-ANP closed subunits (not shown).

The moderate twist of the N2D domain in the BfpD structure is suggestive of a partially open conformation. The N2D twist is similar to that of the pair of subunits in the GmPilB-ANP structure with a N2D twist of 48°, nevertheless with a distinct rotation axis. As Glu338 and the catalytic glutamate Glu295 are 5.3-11.6 Å away from the γ-phosphate of ANP, it appears that the ATPase captured by cryo-EM is a pre-hydrolysis intermediate between the ADP/ATP exchange and ATP hydrolysis steps of the catalysis cycle. The single rotation state of the N2D domain with respect to the CTD in the case of BfpD concurs with its six-fold symmetry.

### Insertions in the BfpD sequence might account for its six-fold symmetry

Amino acid sequences from five type-4 pilus extension ATPase homologues, including three that form two-fold symmetric structures and two from pathogens that have extension ATPases closely related to those of the thermophiles, were compared with BfpD. BfpD showed conservation in the catalytic glutamate, Walker A/B motifs and arginine fingers which are required for nucleotide binding, stability and hydrolysis (Fig. 6). However, several insertions were observed when the N2D and CTD were aligned separately. The N2D contains four conserved arginines, two of which are bound by ATP in the pair of active hydrolyzing subunits in TtPilB (PDB ID: 5IT5). We observed insertions in BfpD around Arg183, Arg217, the linker, and in the catalytic loop that harbors the critical catalytic residue, Glu295 (Fig. 6). We hypothesize that a longer linker and a shift in arginine fingers might be responsible for a different N2D twist seen in BfpD. Furthermore, half of the inter-subunit contacts in BfpD-ANP and BfpD-ADP structures were not conserved among its two-fold symmetric homologues. Taken together, the above sequence differences in the linker and oligomerization interface might be implicated in the observed six-fold symmetry in BfpD.

**Figure 6.**
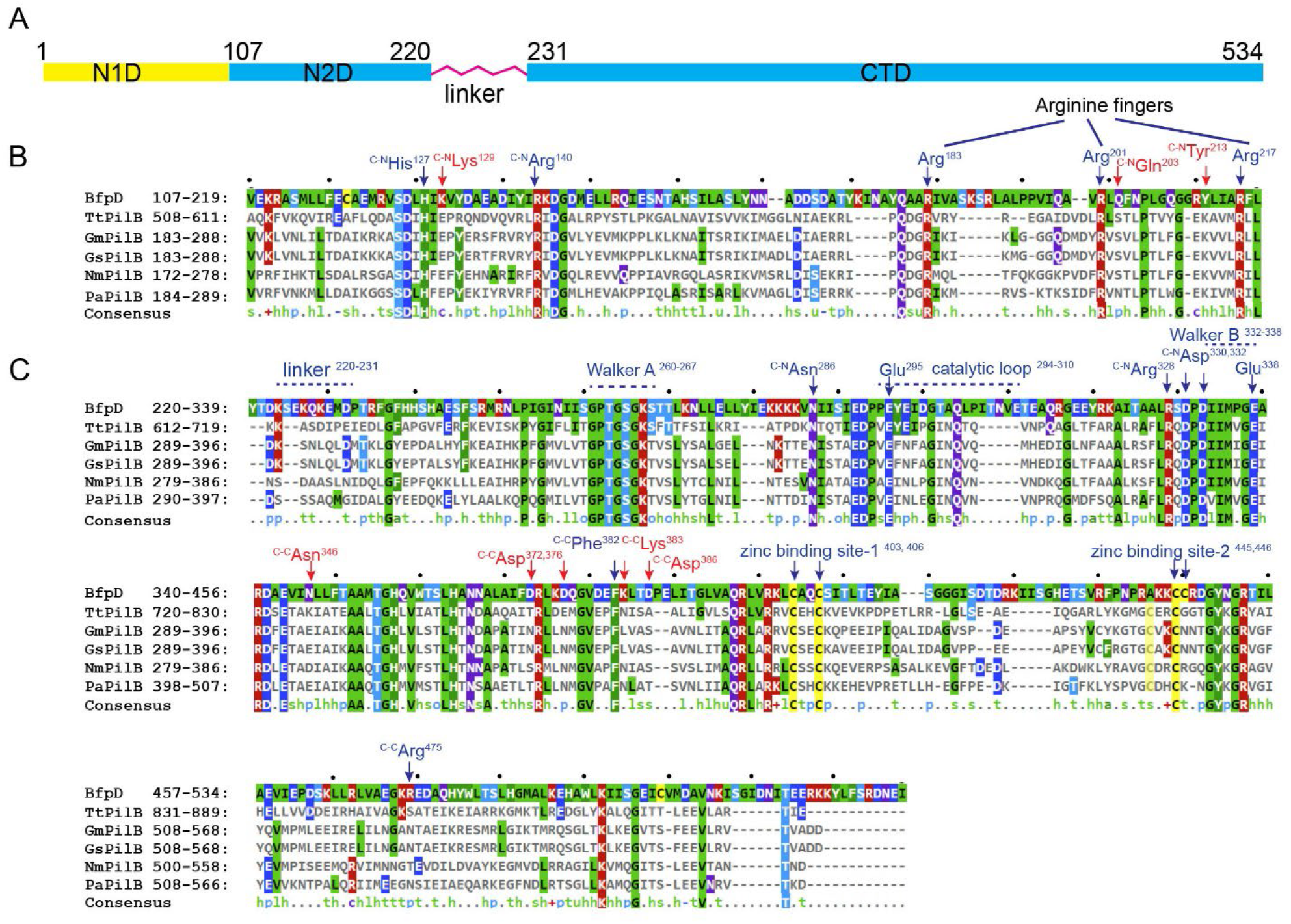
Sequence comparisons of T4P extension ATPase N2D and CTD domains and inter-subunit contacts in BfpD. The alignment of BfpD with five homologues is shown. TtPilB, GmPilB, and GsPilB form two-fold symmetric hexamers and PilB from *P. aeruginosa* and *N. meningitidis* are members of the type-4 pilus ATPases more closely related to them than to BfpD. (A) Domain organization. (B) alignment of the N2D domains shows insertions in BfpD between three conserved Arginine fingers relative to the other enzymes. (C) alignment of the CTD domains shows insertions in the linker region and in the catalytic loop. Sequence elements required for nucleotide binding (Walker A), hydrolysis (Glu295, Glu338) and zinc binding are highlighted. Amino acids forming inter-subunit contacts between CTD and N2D/CTD^+^ of neighboring subunits are indicated, with text specifying the partner contact. Interactions unique to BfpD are colored red.

### BfpD ATPase activity and kinetics

According to molecular modeling with another AAA+-family ATPase (22), the putative catalytic E295 residue in our cryo-EM BfpD model seems to be positioned further from the nucleotide than required for catalysis (Fig. 3 and Movie S2). We therefore examined BfpD activity, using enzyme that had been purified by metal affinity and size exclusion chromatography. We varied the concentration of ATP in the reaction and determined that the apparent *Vmax* and *Km* of BfpD at 0.5 mg/ml are 2.69 ± 0.34 µmole min^-1^ and 239.31 ± 117.63 µM, respectively (Fig. 7A). At an ATP concentration of 3 mM, the specific activity of BfpD was 3.16 ± 0.60 nmol min^-1^ mg^-1^ (Fig 7B), which is consistent with published results for other PilB family extension ATPases (4-7). To assess the role of residues mapped to the catalytic site, we expressed and purified BfpD variants that had mutations in the catalytic glutamate (BfpD_E295C_), and in both that residue and the conserved Walker B glutamate (BfpD_E295C E338Q_). The single and double glutamate mutations reduced the catalytic rate by 5- and 12-fold, supporting their important role in ATP catalysis. However, in each case, we were able to measure specific activity above background spontaneous hydrolysis (Fig 7A). The specific activity of BfpD_E295C_ was 0.69 ± 0.28 nmol min^-1^ mg^-1^ and that of BfpD_E295C E338Q_ was 0.38 ± 0.10 nmol min^-1^ mg^-1^. Importantly, while a plasmid encoding wild type BfpD was able to complement a *bfpD* null mutant to restore bacterial auto-aggregation characteristic of EPEC expressing BFP, plasmids encoding either the E295C or the E338Q BfpD variants were not (Fig. 7C). Thus, the residual activity we were able to detect is insufficient for function.

**Figure 7.**
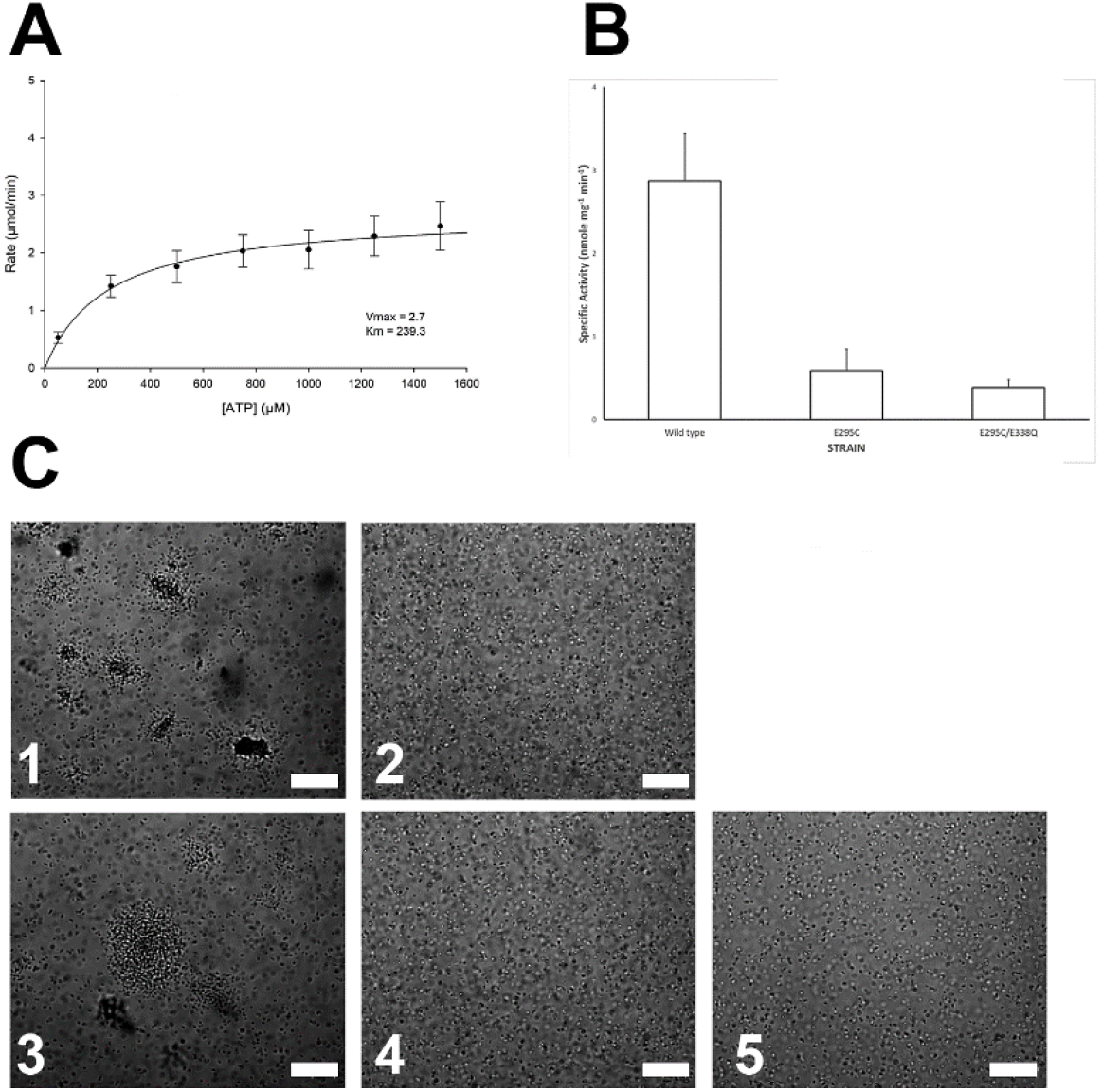
Specific activity, kinetics, and function of BfpD. (A) Rate of inorganic phosphate production as a function of the concentration of ATP for wild type BfpD, from which the apparent *K*_*m*_ and *V*_*max*_ were calculated using Sigma Plot software. Data are from seven biological replicates. (B) Specific activity of BfpD with native (wild type), cysteine substituted for glutamate 295 (single mutant), and both E295C and E338Q substitutions (double mutant). The mean and standard error of the means of seven biological replicates is shown. Analysis of variance revealed significant differences (P < 0.001) between groups. (C) phase-contrast micrographs of (1) wild type, (2) *bfpD* mutant, and *bfpD* mutant strains complemented with plasmids encoding (3) wild type BfpD, (4) BfpD_E295C_, and (5) BfpD_E338Q_. Large aggregates of bacteria indicative of BFP expression are seen in panels 1 and 3. Bars indicate 40 microns.

## DISCUSSION

Using cryo-EM, we determined the structure of BfpD, the extension ATPase of the EPEC bundle-forming T4P. The structure is noteworthy for a number of reasons. It is the first near-atomic structure of a T4P PilB homologue in its native, frozen-hydrated state. It is the first such structure from a pathogenic bacterium. It is an enzyme that is distantly related to its homologues solved to date, all of which were from thermophiles (14). Despite its phylogenetic distance, the BfpD monomer is similar to those of other PilB structures (14-17). As has been the case with all PilB structures solved to date, the full BfpD N-terminus (N1D) was not visualized. In addition, the eight residues intervening the N2D and CTD domains, where flexible linkers exist in the PilB structures, were not resolved here. BfpD was determined either in the presence of ADP or ADP plus ANP. In the BfpD-ANP reconstruction, despite the presence of ADP, ATP and ANP, the six nucleotide binding sites were occupied by the triphosphate or its analogue. Transition from BfpD-ADP to BfpD-ANP resulted in a slight outward shift of the top part of the toroid, a tendency more pronounced in BfpD-ANP class 2. A similar conformational transition was observed by lower-resolution cryo-EM for the TtPilB-ANP structure relative to the structure purified without nucleotide (14, 15). The BfpD monomers are in a nearly identical conformation in both cases. The isolated N2D and CTD domains of BfpD superimpose well with those determined for other PilB structures, while the relative N2D-CTD position is intermediate between the two more extreme open and closed conformations manifested by the other PilB structures.

In contrast to the similarity among monomers of PilB structures, the BfpD quaternary structure revealed unambiguous six-fold symmetry, while those of the PilB enzymes from thermophilic bacteria show two-fold symmetry. The two-fold symmetry is the basis for a symmetric rotary model coupling ATP hydrolysis to large domain shifts, proposed to be translated via the conserved polytopic membrane protein, PilC in the center of the toroid, to lift pilin from the membrane (14, 15). In contrast, a six-fold symmetry is more compatible with a concerted model, in which all catalytic sites act synchronously. Of note, the crystal structure of a catalytically inert C-terminal proteolytic fragment of TtPilB containing the N2D and CTD also displayed two-fold symmetry (16), as does the crystal structure of the N2D-CTD fragment of GsPilB, which was solved in its apo form (17). Thus, ATP hydrolysis is not required for the two-fold symmetry.

TtPilB was also examined by cryo-EM, both bound to AMP-PNP and without exogenous nucleotide, achieving resolution of only ∼8 Å, but, for the first time, the second and third of three predicted N1Ds were observed (16). Comparison of the hexamers in the presence of AMP-PNP and without added nucleotide showed an outward shift in the center of mass of the AMP-PNP structure relative to the structure purified without addition of nucleotide (14, 15). They also observed evidence of displacement of the N1D hexamer by 10-13 Å and suggested an alternative model for energy transduction in which the N1D displacement is linked to pilin through a transmembrane complex of essential T4P biogenesis proteins called PilM, PilN, PilO, and PilP. The BfpD structures in the presence of ADP and AMP-PNP, while not compatible with the symmetric rotary model, show subtle differences in the position of the N2D with respect to the CTD that are similar to those seen in the TtPilB cryo-EM structures (16). These shifts may represent evidence for transduction of mechanical energy at the periphery of the toroid, rather than through PilC at its center. Given that structures of the N1Ds of T4P ATPases have yet to be solved, it is possible that these differences are amplified by conformational changes and transmitted through PilM and PilN to the outer surface of the cytoplasmic membrane. Support for this alternative model is found in the structure of the related type 2 secretion ATPase N1D in complex with the PilM homologue (23) and in the complex formed by a T4P pilin with the PilM, PilN, PilO transmembrane assembly (24).

The conformation of the BfpD nucleotide binding site raises interesting questions about its catalytic activity, as the distances of the putative catalytic Glu295 and of the conserved Walker B Glu338 are approximately 5.3-11.6 Å and 6 Å from the phosphorus atom in the ANP γ-phosphate, respectively. This appears to be too far for activation of the water molecule that is responsible for hydrolysis (20). By comparison, the corresponding distances of 6-6.5 Å at the tightest interface in the GmPilB-ANP structure, suggesting that all subunits in the BfpD-ANP structure are in a pre-hydrolytic conformation. Nevertheless, we were able to demonstrate that BfpD has ATPase activity that is comparable to that reported for several other such enzymes (4-8). Surprisingly, we were also able to measure ATPase activity above background in a BfpD variant that has mutations in both Glu295 and Glu338, which suggests that mere binding of ATP to the open pocket facilitates hydrolysis. However, the residual activity is insufficient for pilus biogenesis, as demonstrated by expressing these variants in a *bfpD* null mutant. We further suggest that neither the *in vitro* activity we were able to measure in wild type BfpD, nor that of any other PilB homologue is sufficient to power T4P extension. BfpD specific activity of about 3 nmol min^-1^ mg^-1^ corresponds to 0.02 ATP molecules hydrolyzed per second per hexamer. In contrast, given an axial rise of 10.5 Å, corresponding to 952 subunits per micron in a T4P (25), real-time video demonstrates that the rate of T4P extension equates to 333 - 1072 subunits added per second (10-12). Assuming that each subunit added requires at least one ATP, these *in vitro* enzyme activity measurements fall short of the values required *in vivo* by more that 10,000-fold. If six ATPs are required per subunit, this shortfall multiplies correspondingly. Modifications to the *in vitro* assays, such as addition of phospholipids or partner proteins (26), or imposed hexamerization (27), as well adjustments to the assumptions, would do little to alter this arithmetic. A similar conclusion was reached by the authors of the study that described a thermophilic PilB enzyme with specific activity of 700 nanomoles mg^-1^ min^-1^ (9). They suggested that enzyme activity must be higher *in vivo* where partner proteins are in optimal orientation and concentration. Interestingly, these authors described complex enzyme kinetics including substrate inhibition at concentrations exceeding 1.5 mM ATP and a non-linear relationship between substrate concentration and catalytic rate below that concentration. They interpreted these results as consistent with the symmetric rotary model of catalysis that had been proposed for TtPilB. In contrast, BfpD displayed simple Michaelis-Menton kinetics compatible with a concerted model. The fact that all enzyme subunits were occupied by the nucleotide triphosphate or triphosphate analogue in the presence of ADP, ATP, and ANP is also consistent with a concerted model.

In summary, we report the first 3D structure of a T4P extension ATPase from a human pathogen. The BfpD structure determined under cryogenic conditions in a native state, unparalleled in its detail, in the presence of ADP and a non-hydrolysable ATP homologue, lays a structural foundation to understand similarities and differences of T4P machinery among different clades. The study expands our understanding of mechanisms of catalysis, activation, and energy transduction of the PilB family of T4P extension ATPases.

## METHODS

### Site-directed mutation of putative BfpD active site residues

All bacterial strains and plasmids used in this study are listed in Table S2. A codon-optimized *bfpD* gene in plasmid BfpD-Hcp (GenScript, USA, a kind gift from Dr. Kurt Piepenbrink, Fig. S5) was amplified using PCR with primers BfpDNcoI and BfpDXhoI (Table S3), creating a Leu to Val substitution at codon two to allow cloning into a pET30a plasmid vector at NcoI/XhoI sites and creating plasmid pJZM005. Fast-Cloning (28) was used to introduce substitutions for glutamate codons at amino acid positions 295 and 338 of BfpD in plasmids pJZM005 and pJZM032, respectively. In short, a pair of primers (Table S3) was designed such that each has a complementary sequence including the mutated codon and divergent sequences overlapping with *bfpD*. Codons specifying neutrally charged hydrophilic amino acids of similar size to glutamate were chosen to minimize disruptions to structure and those that could be expressed and purified were studied further. The PCR products were digested with DpnI and subsequently transformed into *E. coli* DH5α competent cells. The plasmids pJZM032 and pJZM042 encoding BfpDE295C and BfpD_E295C E338Q_ respectively, were confirmed by sequencing, expressed, and purified from *E. coli* BL21(DE3) as described below. To make plasmids for *in vivo* complementation, we first constructed a plasmid harboring wild type *bfpD* in low-copy-number cloning vector pWKS30 (29). To do this, *bfpD* with its N-terminal His tag and S tag, was isolated from pRPA405 (30) as an XbaI-SacI fragment, and inserted into pWKS30. The resultant plasmid pEMM1 was later discovered to lack its native stop codon, while a stop codon on the vector was noted downstream, thus adding an elongated non-native C-terminus to the predicted protein. We used FastCloning to restore a TAG stop codon in its original position in pEMM1, and the new plasmid was named pJZM031. Thereafter, pJZM031 was used as template to introduce E295C and E338Q mutations to obtain pJZM032 and pJZM036, respectively. FastCloning was used to introduce the E295C mutation. For the E338Q mutation, we first introduced the mutation using overlap extension PCR (31), which was later cloned into pJZM031 (XbaI/SacI digested). Auto-aggregation and disaggregation were assessed as previously described (30). Briefly, overnight cultures of E2348/69, UMD926 and VCU019 containing pWKS30, pJZM031, pJZM032, or pJZM036 were diluted 100-fold in Dulbecco’s modified Eagle’s medium (Corning) and grown for 4 h at 37 °C before examination by phase-contrast microscopy.

### BfpD, BfpD_E295C_ and BfpD_E295C E338Q_ expression and purification

For purification of BfpD, *E. coli* strain BL21(DE3) pJZM005 was grown at 37°C in Luria-Bertani medium to an optical density at 600 nm (OD_600_) of 0.6 and induced with 1 mM isopropyl β-D-1-thiogalactopyranoside (IPTG) at 16°C overnight. Cells were harvested by centrifugation, sonicated in lysis buffer (50 mM PO_4_, 300 mM NaCl, pH 8.0, 10 mM imidazole), and purified by affinity chromatography on Cobalt-NTA resin, created by stripping nickel from a Ni-NTA (Qiagen, USA) column with 100 mM EDTA and replacing with 10 mM CoCl_2_. Fractions eluted with 250 mM imidazole were analyzed by SDS page, combined, and dialyzed against buffer (20 mM Tris–HCl, pH 7.6, 100 mM NaCl, 1 mM MgCl_2,_ 2 mM DTT) at 4°C. For some experiments, a Suprose6 10/300 column was used to achieve further purification, as noted.

### Negative staining and 2D averaging

BfpD in 20 mM Tris (pH 7.6), 0.1 M NaCl, 5 mM MgCl_2_, 2 mM DTT and 1 mM ADP was diluted to 0.01-0.03 mg/ml and stained with 0.75% uranyl nitrate using an established protocol (32). Images were acquired at 50000X magnification in low dose mode with a Tecnai F20 microscope operated at 120kV. Reference-free 2D averages were obtained with the SPIDER program (33).

### Cryo-EM grid preparation and data acquisition

The cryo-EM sample buffer consisted of 20 mM Tris (pH 7.6), 0.1 mM NaCl, 5 mM MgCl_2_, 2mM DTT and 0.5 mM CHAPSO, and additionally either a mixture of ADP (1 mM), ATP and ANP at a 2:4:5 millimolar ratio or 1 mM ADP alone. Usage of CHAPSO improved BfpD orientation in vitreous ice which enabled successful 3D reconstruction. The 300 mesh UltraAufoil -1.2/1.3 holey-gold (Quantifoil, Germany) grids were cleaned with a customized protocol (34) prior to glow discharge. Purified BfpD was diluted to 4.5 mg/ml and 3 µl was applied to a glow-discharged 300 mesh UltraAufoil -1.2/1.3 holey-gold (Quantifoil, Germany). Grids were blotted for 2 s with ash-free Whatman^®^ Grade 540 filter paper in a Vitrobot Mark IV (ThermoFisher Scientific) and plunged into liquid ethane. Sample quality and distribution was assessed on a Tecnai F20 (ThermoFisher Scientific) electron microscope. Data acquisition was carried out in a Titan Krios transmission electron microscope (ThermoFisher Scientific) operated at 300 kV and counting mode, with a Gatan K3 detector and a 10 eV slit width Gatan Quantum Energy Filter (GIF). Datasets were collected in automated mode with the program Latitude (Gatan) with cumulative electron dose of 60 e-/Å^2^ applied over 40 frames.

### Single-particle image processing

Movie stacks collected for BfpD-ANP and BfpD-ADP datasets were processed in cryosparc2.15. Gain-normalization, movie-frame alignment, dose-weighting, and full and local motion correction were carried out with the patch motion correction. Global and local contrast transfer function values were estimated from non-dose weighted motion-corrected images using patch CTF module. Bad micrographs with ice and ethane contamination and poor CTF fits were discarded. Subsequent image processing operations were carried out using dose-weighted, motion-corrected micrographs. 2D-class average images obtained from 1000 manually picked particles were used to pick about 2.83 and 1.04 million particles in ANP and ADP datasets. Extensive 2D-classifications of 4-8 rounds yielded 424708 and 313223 pure particles which led to 2.98 and 3.1 Å consensus maps. The reported resolutions of the cryo-EM maps are based on FSC 0.143 criterion (35). Six-fold rotational symmetry (C6) confirmed from the 2D class averages was applied during 3D refinement. The density of ANP was confirmed at the interphase of N2D and CTD, though 2.5:1 millimolar mixture of ANP and ADP was used in the mixed nucleotide dataset. A summary of image processing of the ANP and ADP datasets can be found in Figs S1, S2, respectively.

### *De novo* model building and structure refinement

A crude model for a single BfpD subunit was built using the ADP map with *Phenix*.*map_to_model* tool (36), which was improved by chain tracing. The regions from 107-223 (N2D), 231-534 (CTD) were built. The density for the N-terminus (N1D; 1-106) and flexible linker connecting N2D and CTD (224-230) was not visualized. A clear density for a loop spanning 294-310 in the ADP structure was obtained which is disordered in the ANP structure. The monomeric model was expanded to a hexamer model. Local density fit of the modeled sequence was improved over an iterative process of amino acid fitting in Coot (37) alternated with real space refinement in PHENIX (38). Real space refinement was carried out with NCS constraints and secondary structure and Ramachandran restraints. Comprehensive model validation was carried out with PHENIX and PDB validation server at https://validate-rcsb-2.wwpdb.org/ and are summarized in Table S1. Surface charge was calculated with chimera (39). Figures were generated with The PyMOL Molecular Graphics System, Version 2.0 Schrödinger, LLC (https://pymol.org) and chimera.

### ATPase activity

ATPase activity was measured using a previously established method (40, 41) with slight modifications. In brief, stocks of BfpD purified by cobalt affinity chromatography were prepared in assay buffer (150 mM Tris–HCl, pH 7.6, 300 mM NaCl, 1 mM MgCl_2,_ 2 mM DTT) to achieve 0.5 mg/ml in the final reaction and mixed with various concentrations (50 µM - 1500 µM) of freshly prepared ATP in the same assay buffer. The reaction mixtures (in triplicate) were aliquoted to 96-well plates (one plate for each time point) and incubated at 37°C. At the defined time points, the reaction was stopped by adding 100 µl of the assay reagent (a 3:1 mixture of freshly prepared 0.045% malachite green hydrochloride in water and 4.2% ammonium molybdate in 4 N HCl, along with 1% Triton X-100) and followed by 20 µL of citrate solution (34%). The absorbance at 655 nm was measured on a Clariostar Monochromator Microplate Reader (BMG Labtech). The ATPase activity was extrapolated from a standard curve of a set of defined phosphate concentrations present in two columns of each plate. To measure apparent *K*_*m*_ and *V*_*max*_, various ATP concentrations, and the reaction was terminated at multiple time points. Released PO_4_ concentration was plotted as a function of time for the different ATP concentrations and the slope was used as initial velocity (*V*_*0*_). Data from seven biological replicates were fit by nonlinear regression to the Michaelis-Menten equation with SigmaPlot® (Systat Software, Inc.) to calculate apparent *K*_*m*_ and *V*_*max*_. The specific activity of BfpD was measured by varying BfpD concentrations (0.125 – 2.0 mg/ml) in the assay buffer (above) with 3 mM ATP and a 30 min reaction time at 37°C. The rate of T4P extension in microns per time was estimated from several studies with various methods (10-12) and from an estimate of 952 T4P subunits per micron in the pilus (25).

## Appendix

Supplementary material for this article is available in a separate file.

## Data availability

Cryo-EM maps of BfpD-ANP class-1, 2 and BfpD-ADP have been deposited in the Electron Microscopy Databank (EMDB) with accession codes 27795, 27796 and 27797 respectively. Atomic models of BfpD-ANP class-1, 2 and BfpD-ADP have been deposited in the RCSB PDB database with accession codes 8DZE, 8DZF, and 8DZG respectively.

## Acknowledgments

We are grateful to Kurt Piepenbrink (currently at University of Nebraska) for a providing codon-optimized BfpD construct. Cryo-grid preparation and screening were carried out at the Cryo-EM Unit at Virginia Commonwealth University (VCU), and data collection was carried out at the Molecular Electron Microscopy Core Facility at the University of Virginia (UVA) (supported by NIH U24 GM116790). We thank Dr. Kelly Dryden for cryo-EM data collection. Supported by VCU Accelerate Fund (to M.D. and M.S), NIH R56 AI111767 (to M.D.), and NIH R01 AR068431 (to M.S).

## Author contributions

J.L. performed protein purification and mutagenesis; P.K.S. performed enzyme assays; A.R.N. performed protein cryo-grid preparation, image processing and model building; A.R.N, M.S. and M.D. interpreted the structural data; M.D. and M.S conceived, designed and supervised all experiments, M.D., M.S., and A.N. wrote the manuscript.

