## Supplemental Figures and Tables for "Cryo-EM structure of the Type IV pilus extension ATPase from enteropathogenic *Escherichia coli*"

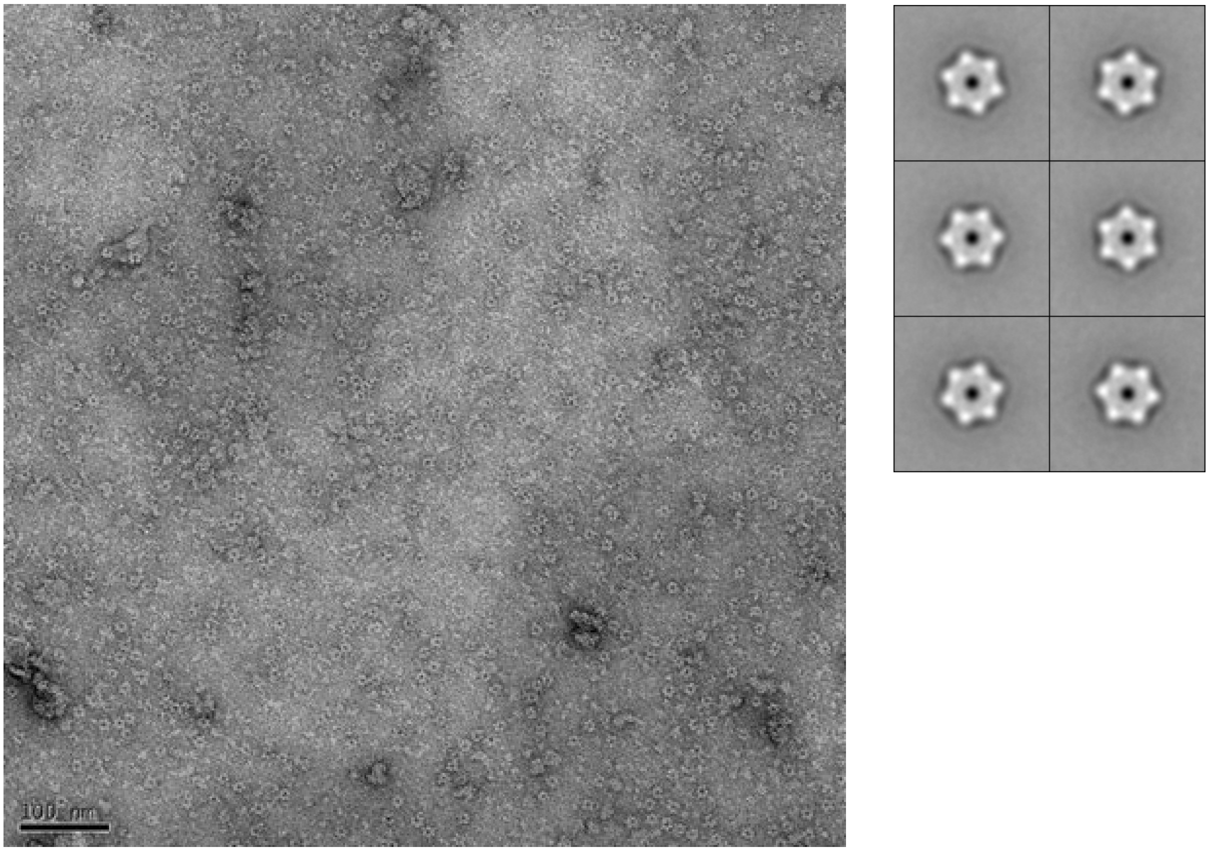


**Figure S1.** BfpD shows six-fold rotational symmetry

Left, Negative stained image of BfpD stained with 0.75% uranyl formate and displayed at 50,000x magnification. The image was acquired in low dose mode at 120kV using a Tecnai F20 microscope. Right, representative 2D averages.

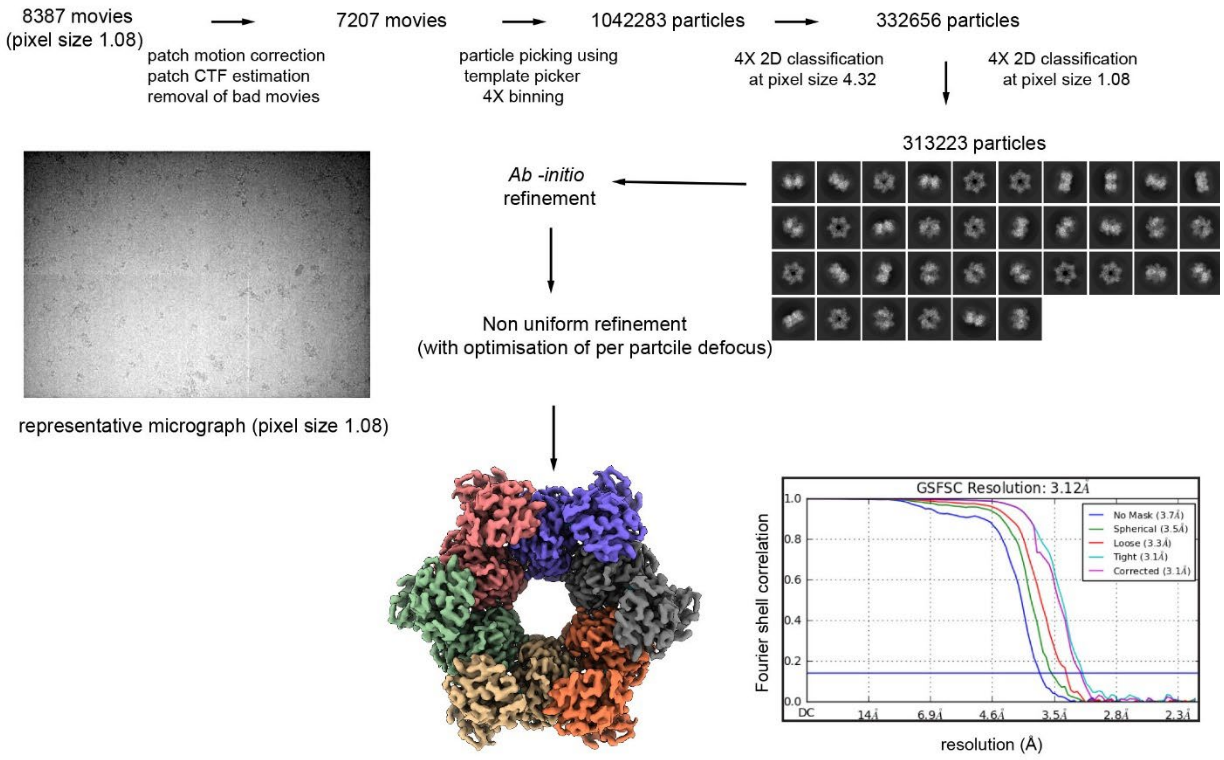


**Figure S2.** Image processing of the BfpD-ADP dataset in cryoSPARC 2.15.

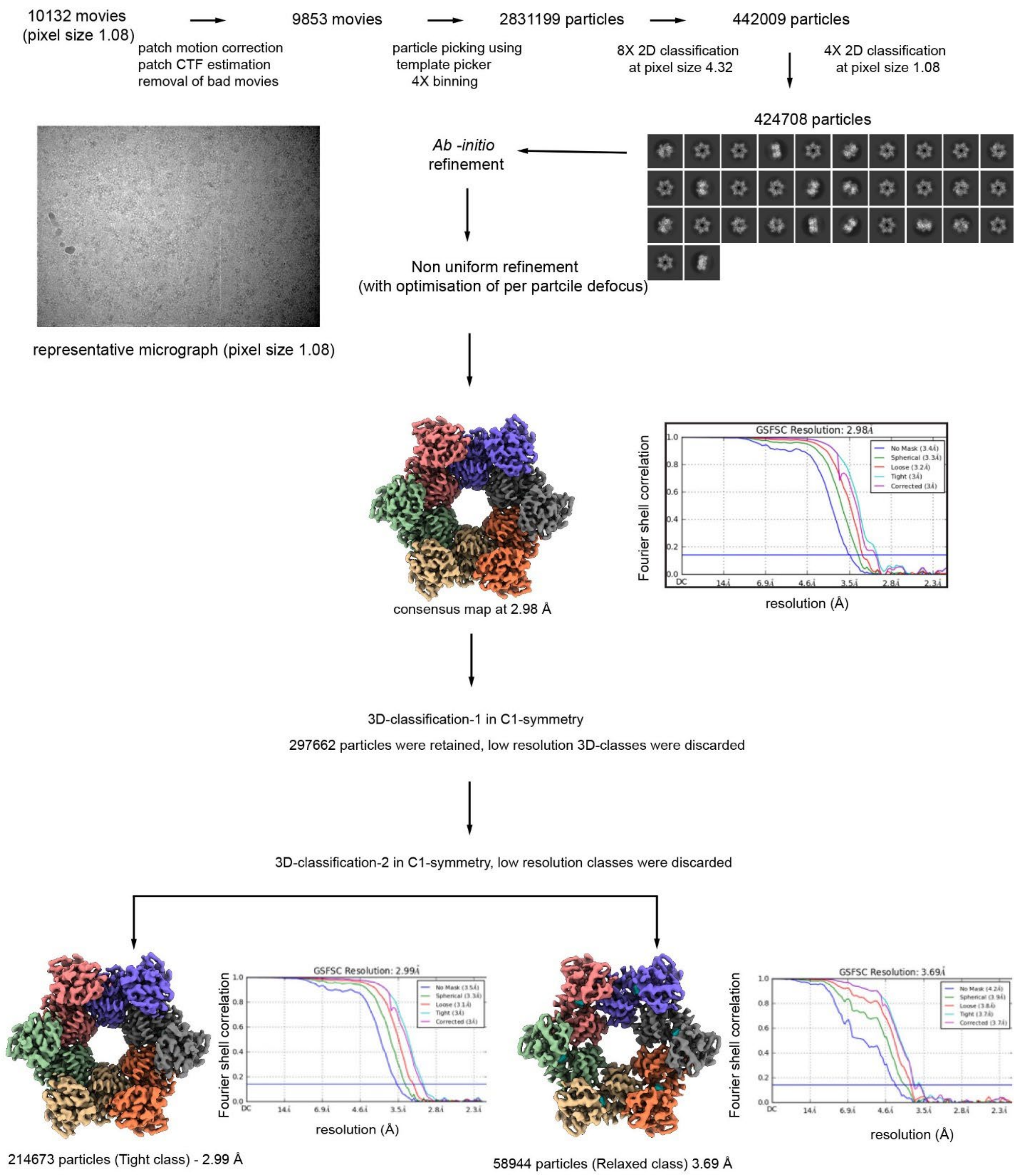


**Figure S3.** Image processing of the BfpD-ANP dataset in cryoSPARC 2.15.

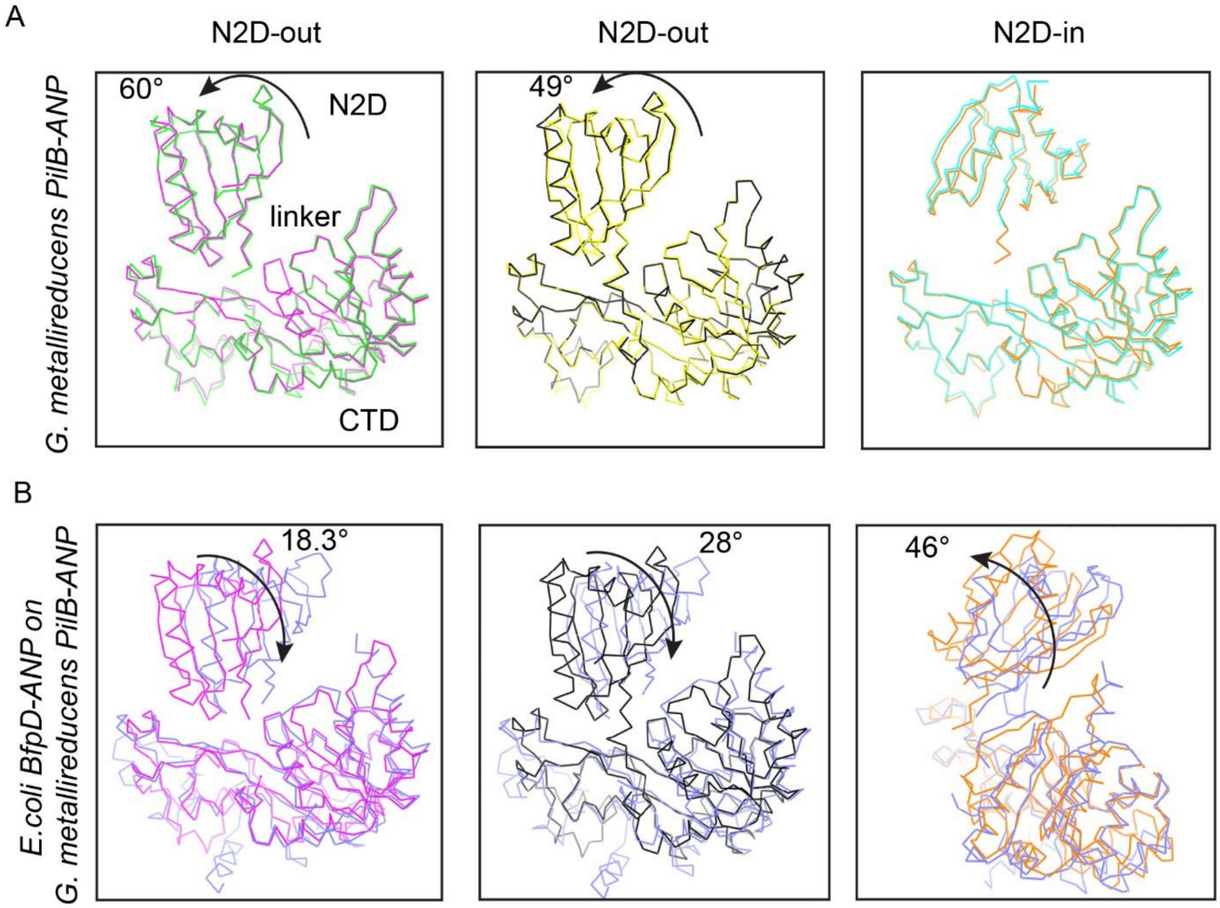


**Figure S4.** The intermediate open subunits of BfpD differ from *Geobacter metallireducens* PilB subunits

A, From left, Superposition of pairs of opposite subunits from *Geobacter metallireducens (Gm)* PilB-ANP crystal structure (pdb id: 5TSH). The N2D domain exists in three different rotation states; the two N2D-out conformations are rotated by 49° and 60°, away from the center of the elongated hexamer, when compared to the N2D-in conformation. B, From left, Comparison of the N2D domains of BfpD-ANP (in blue) and GmPilB-ANP N2D conformations, with the C-terminal domains superimposed. Out of plane rotations of 18° and 28° are required to attain BfpD-ANP N2D conformation. N2D domain of BfpD rotated ~46° from the N2D-in conformation from GmPilB structure. The structures in the right are rotated in-plane for clarity. The plane of rotation in BfpD-ANP N2D in B differs from the inter-subunit comparison in GmPilB-ANP structure in A.

ATGCACCATCATCATCATCATTCTTCTGGTCTGGTGCCACGCGGTTCTGGTATGAA
AGAAACCGCTGCTGCTAAATTCGAACGCCAGCACATGGACAGCCCAGATCTGGGT
ACCGACGACGACGACAAGGCCATGGTGAACAAAACCGAAAAAACCAGCGATCTGA
TGTTTGAACGCTTTAAACGTAACGTGAGCGAAATTGTTACCGGTGATGGTGGTGAA
CTGGAACTGACCGTTGAACAGCGTAAATACTTTCTGATCTTCAAAAACGGCGATTTT
CTGGTTAGCAGCTGCCATATGAAACATCATCTGGTTCAGATGCTGCGTGAAATTGC
AACCCGTAAAGGTTATCCGAATCTGACCATTTATGAGGTGAACCTGAAAGATATTC
GCCTGCTGTATGAAGCAAGCCTGAAAACCGTTCAGAATAATGGTCAGGATCTGCTG
CCGGTTGAAAAACGTGCAAGCATGCTGCTGTTTGAATGTGCAGAAATGCGTGTTAG
CGATCTGCATATCAAAGTTTATGATGCAGAGGCCGATATCTATATTCGCAAAGATG
GTGATATGGAACTGCTGCGTCAGATTGAAAGCAATACCGCACATAGCATTCTGGCA
AGCCTGTATAATAACGCAGATGATAGTGATGCCACCTACAAAATTAACGCATATCA
GGCAGCACGTATTGTTGCAAGCAAAAGCCGTCTGGCACTGCCTCCGGTTATTCAG
GCAGTTCGTCTGCAGTTTAATCCGCTGGGTCAGGGTGGTCGTTATCTGATTGCCC
GTTTTCTGTATACCGATAAAAGCGAGAAACAGAAAGAAATGGATCCGACCCGTTTT
GGTTTTCATCATAGCCATGCAGAAAGCTTTAGCCGTATGCGTAATCTGCCGATTGG
CATTAACATTATTAGCGGTCCGACCGGTAGCGGTAAAAGCACCACACTGAAAAATC
TGCTGGAACTGCTGTACATCGAGAAAAAAAAAAAAAGTGAACATCATCAGCATCGAG
GATCCGCCTGAATATGAAATTGATGGCACCGCACAGCTGCCGATTACCAATGTTGA
AACCGAAGCACAGCGTGGTGAAGAATATCGTAAAGCCATTACCGCAGCACTGCGT
AGCGATCCGGATATCATTATGCCTGGTGAAGCACGTGATGCCGAAGTTATTAATCT
GCTGTTTACAGCAGCAATGACAGGTCATCAGGTTTGGACCAGCCTGCATGCAAATA
ATGCACTGGCAATTTTTGATCGTCTGAAAGATCAGGGTGTGGATGAATTTAAACTG
ACCGATCCGGAACTGATTACCGGTCTGGTTGCACAGCGTCTGGTTCGTAAACTGT
GTGCACAGTGTAGCATTACCCTGACCGAATATATTGCAAGCGGTGGTGGTATTAGC
GATACCGATCGTAAAATCATTAGTGGTCATGAAACCAGCGTTCGTTTTCCGAATCC
GCGTGCAAAAAAATGTTGTCGTGATGGTTATAATGNTCGTACCATTCTGGCGGAAG
TTATTGAACCGGATAGCAAACTGCTGCGCCTGGTTGCCGAAGGTAAACGTGAAGA
TGCACAGCATTATTGGCTGACCTCACTGCATGGTATGGCACTGAAAGAACATGCAT
GGCTGAAAATCATTTCAGGCGAAATTTGTGTTATGGATGCCGTCAACAAAATTAGC
GGCATTGATAACATTACCGAAGAACGCAAAAAATACCTGTTCAGCCGTGATAATGA
AATTTAG

**Figure S5.** Codon optimized *bfpD*. Underlined sequence is from pET30a encoding the start codon and hexa-histidine tag.

**Movie S1.** Concerted N2D domain movement in BfpD during ATP binding and hydrolysis

Top and side view; As a result of N2D twisting in each of the six subunits, the BfpD ring contracts during the transition from ANP class-2 to ADP and vice versa. The ring closure represents an event of ATP-bound BfpD, progressing to the post ATP hydrolysis state (ADP). In the active hydrolyzing state, the N2D-CTD of BfpD is expected to have a more compact conformation than the states depicted here.

**Movie S2.** Transition in the BfpD active site from ANP-class-2 to ANP-class-1

The N2D moves toward the six-fold axis in going from ANP class-2 to ANP-class-1, likely reflecting two successive steps on the way to ATP hydrolysis. The ring contraction from class-2 to class-1 is accompanied by loss of contact between the ATP-stabilizing Arg217 and the ɣ phosphate of ATP, and gradual shortening of the distance between the catalytic residue Glu295 and that same phosphate, priming the active site for catalysis.

**Table S1.** Summary of cryo-EM image processing and model building statistics

<table>
<tr><th></th><th>BfpD-ADP</th><th>BfpD-ANP (class-1)</th><th>BfpD-ANP (class-2)</th></tr>
<tr><td>Data Acquisition</td><td></td><td></td><td></td></tr>
<tr><td>Microscope/Detector</td><td>Krios/K3</td><td colspan="2">Krios/K3</td></tr>
<tr><td>Voltage (kV)</td><td>300</td><td colspan="2">300</td></tr>
<tr><td>Magnification</td><td>81000</td><td colspan="2">81000</td></tr>
<tr><td>Data collection mode</td><td>Counting</td><td colspan="2">Counting</td></tr>
<tr><td>Pixel Size (Å) (super-resolution)</td><td>1.08</td><td colspan="2">1.08</td></tr>
<tr><td>Focus range (μm)</td><td>-1. to -2.25</td><td colspan="2">-1. to -2</td></tr>
<tr><td>Total electron dose (e/Å$^2$) (Number of frames)</td><td>60 (40)</td><td colspan="2">60 (40)</td></tr>
<tr><td>Total number of movies</td><td>8387</td><td colspan="2">10132</td></tr>
<tr><td>Image Processing</td><td></td><td></td><td></td></tr>
<tr><td>Total number of particles picked</td><td>1042283</td><td colspan="2">2831199</td></tr>
<tr><td>Particles after 2D classification</td><td>332656</td><td colspan="2">442009</td></tr>
<tr><td>Particles used for 3D refinement</td><td>313223</td><td>214673</td><td>68944</td></tr>
<tr><td>Resolution (Å)</td><td>3.12</td><td>2.99</td><td>3.69</td></tr>
<tr><td>EMDB ID</td><td>27797</td><td>27795</td><td>27796</td></tr>
<tr><td>Model Refinement</td><td></td><td></td><td></td></tr>
<tr><td>RMS Deviation (Bonds)</td><td>0.003</td><td>0.003</td><td>0.004</td></tr>
<tr><td>RMS Deviation (Angle)</td><td>0.712</td><td>0.647</td><td>0.892</td></tr>
<tr><td>Ramachandran Plot statistics (%)</td><td></td><td></td><td></td></tr>
<tr><td>Preferred</td><td>95.39</td><td>92.84</td><td>93.58</td></tr>
<tr><td>Allowed</td><td>4.37</td><td>6.91</td><td>6.17</td></tr>
<tr><td>Outliers</td><td>0.24</td><td>0.25</td><td>0.25</td></tr>
<tr><td>Model Validation</td><td></td><td></td><td></td></tr>
<tr><td>Clash-score</td><td>2.36</td><td>4.94</td><td>6.51</td></tr>
<tr><td>MolProbity Score</td><td>1.38</td><td>1.71</td><td>1.78</td></tr>
<tr><td>EMRinger score</td><td>3.29</td><td>2.86</td><td>1.35</td></tr>
<tr><td>PDB ID</td><td>8DZG</td><td>8DZE</td><td>8DZF</td></tr>
</table>

**Table S2.** Strains and plasmids used in this study

| Strain or Plasmid | Genotype or description | Source or reference |
|---|---|---|
| **Strains** | | |
| DH5α | *supE44 ΔlacU169(φ80 lacZΔM15) hsdR17 recA1 endA1 gyrA96 thi-1 relA1* | Invitrogen |
| BL21(DE3) | $F^{-}$ *omp*T *hsd*$S_B$ ($r_B^{-}$, $m_B^{-}$) *gal dcm* (DE3) | Invitrogen |
| E2348/69 | Nalidixic acid-resistant variant of prototypic virulent O127:H6 clinical EPEC isolate | (1) |
| UMD926 | E2348/69 *bfpD::aphA3* | (2) |
| **Plasmids** | | |
| pET30a | Expression plasmid; $Km^r$ | Novagen |
| BfpD-Hcp1 | Contains codon optimized *bfpD* gene fused to Hcp1 | Kurt Piepenbrink |
| pJZM005 | Codon-optimized *bfpD* in pET30a | This study |
| pJZM032 | pJZM031 with $BfpD_{E295C}$ | This study |
| pJZM042 | pJZM031 with $BfpD_{E295C,\ E338Q}$ | This study |
| pWKS30 | Low copy number vector; $Ap^r$ | (3) |
| pRPA405 | Plasmid having *bfpD* gene with N-terminus His tag and S tag | (4) |
| pEMM1 | *bfpD* subcloned into pWKS30 but lacking its native stop codon. | This study |
| pJZM031 | Wild type *bfpD* complementation plasmid with corrected stop codon from pEMM1 | This study |
| pJZM032 | pJZM031 with $BfpD_{E295C}$ | This study |
| pJZM036 | pJZM031 with $BfpD_{E338Q}$ | This study |

**Table S3.** Primers used in this study.

| **Name** | **Sequence* (5` - 3`)** | **Target** | **Purpose** |
|---|---|---|---|
| BfpDNcoI | TTGATCCCTTCA**CCATGG**TG | *bfpD* | Amplification of *codon optimized bfpD gene* |
| BfpDXhoI | TGACGT**CTCGAG**ctaaatttcattatcacggctgaac | *bfpD* | |
| BfpD E295C.P1 | gatccgcct**tgc**tatgaaattgatggcaccgcac | *bfpD* | *For creation of bfpD*$_{E295C}$ |
| BfpD E295C.P2 | aatttcata**gca**aggcggatcctcgatgctg | *bfpD* | |
| E338Q.P1 | cacgtgcc**ctg**accaggcataatgatatccgg | *bfpD* | *For creation of bfpD*$_{E338Q}$ |
| E338Q.P2 | tgcctggt**cag**gcacgtgatgccgaagttat | *bfpD* | |
| pEMM1TAG.P1 | aacgagataTAGGAATTCGAGCTCCAATTCG | *bfpD* | Stop codon TAG insertion in pEMM1 Making plasmid pJZM031 |
| pEMM1TAG.P2 | GAGCTCGAATTCCTAtatctcgttatctctgctg | *bfpD* | |
| wtbfpDE295C.P1 | tcaatttcatagcaaggtggatcttcaatgctg | *bfpD* | *bfpD*$_{E295C}$ for complementation. Making plasmid pJZM032 |
| wtbfpDE295C.P2 | atccaccttgctatgaaattgacggcacggc | *bfpD* | |
| wtbfpDE338Q.P11 | AGCTTGATATCGAATTCCTGC | *bfpD* | |
| wtbfpDE338Q.P2 | ccctggcctgccctggcattattatatccgg | *bfpD* | *bfpD*$_{E338Q}$ for complementation. Making plasmid pJZM036 |
| wtbfpDE338Q.P1 | gccagggcaggccagggatgctgaagtga | *bfpD* | |
| wtbfpDE338Q.P22 | ACTATAGGGCGAATTGGAGC | *bfpD* | |

*Nucleotides underlined and with bold font indicate restriction sites or mutated codons. Gene coding nucleotides are indicated by lowercase letters, while uppercase letters highlight plasmid based or additional exogenous nucleotides added to facilitate cloning of the specified gene.

1. Nisa, S., Hazen, T. H., Assatourian, L., Nougayrede, J. P., Rasko, D. A., and Donnenberg, M. S. (2013) In vitro evolution of an archetypal enteropathogenic *Escherichia coli* strain. *J Bacteriol* **195**, 4476-4483

2. Anantha, R. P., Stone, K. D., and Donnenberg, M. S. (2000) Effects of *bfp* mutations on biogenesis of functional enteropathogenic *Escherichia coli* type IV pili. *J Bacteriol* **182**, 2498-2506

3. Wang, R. F., and Kushner, S. R. (1991) Construction of versatile low-copy-number vectors for cloning, sequencing and gene expression in *Escherichia coli*. *Gene* **100**, 195-199

4. Crowther, L. J., Anantha, R. P., and Donnenberg, M. S. (2004) The inner membrane subassembly of the enteropathogenic Escherichia coli bundle-forming pilus machine. *Mol Microbiol* **52**, 67-79
